# Small-scale within-drainage spawning behavior causes population differentiation in Atlantic salmon

**DOI:** 10.64898/2026.07.03.736392

**Authors:** Flavio Di Giorgio, Cintia Oliveira Carvalho, Johanna Sjöstedt, Martin I. Lind, Raphael Gollnisch, Anders Persson, Olle Calles, Samuel Shry, P. Anders Nilsson

## Abstract

Understanding the genetic structure of keystone species within river networks is essential for effective conservation and management. While population differentiation of anadromous species often occurs between river systems, less research has been conducted on differentiation within rivers with smaller catchment areas. In this study, we investigated the population genetic structure of wild Atlantic salmon (Salmo salar) across the small-scale river Rönne å system in southernmost Sweden using Restriction-site Associated DNA sequencing (RADseq). Although the Admixture analysis did not detect clearly defined genetic clusters, significant pairwise FST values and DAPC revealed emerging population differentiation among the Rönne å tributaries. The observed patterns are consistent with a system characterized by connectivity, where genetic flow is present but can be reduced by behavioral and ecological factors such as spawning homing behavior and selective movements. These findings suggest that, despite overall connectivity, Atlantic salmon populations in the Rönne å catchment area may function as partially independent sub-populations. This highlights the importance of conservation and management strategies in fragmented river systems to consider population genetic structure to support resilient salmon populations under ongoing anthropogenic pressures.

## Introduction

Genetic differentiation may arise from different forms of isolation acting within and/or between species and populations (Westram et al., 2022). When populations experience contrasting environmental conditions, divergent selection can promote local adaptation, leading to differences in e.g. morphology, physiology, behavior, and life-history traits that reflect adaptive responses to selective pressures (Wadgymar et al., 2022). For instance, high genetic diversity within populations has been shown to enhance ecosystem processes and resilience, with effects on productivity, nutrient cycling, and stability (Fargeot et al., 2025). Conversely, loss of genetic variation can reduce adaptive potential, diminish phenotypic diversity, and weaken responses to environmental stressors, increasing extinction risk and undermining ecosystem services (Kardos et al., 2021). Failure to recognize genetically distinct populations or evolutionary significant units may lead to inappropriate management actions (Segelbacher et al., 2022). Therefore, conserving genetic diversity is not only central to preserving evolutionary processes, but also to maintain ecosystem functioning, and the long-term effectiveness of conservation strategies under global change.

While much attention has been given to spatial and temporal isolation, behavioral isolation can be equally influential in driving differentiation (Mendelson and Safran, 2021). Behavioral isolation occurs when individuals preferentially mate with members of their own population or species, due to differences in mating behaviors, such as distinct courtship songs, dances, or signals, which promote recognition of potential mates and thus reduce or eliminate interbreeding between groups (Barerra et al., 2024). In sticklebacks (*Gasterosteus aculeatus*), for example, males exhibit distinctive courtship behaviors and nuptial coloration that females of other populations rarely recognize, leading to reproductive isolation (McKinnon and Rundle, 2002). The females of Bahamas mosquitofish (*Gambusia hubbsi*) have also shown resistance toward foreign males, expressed through aggression and reduced mating success, representing a mechanism that contributes to behavioral isolation (Pärssinen et al., 2026). Moreover, in salmonids, natal homing behavior results in individuals returning to their birth streams to spawn, creating spatial genetic structuring and limiting gene flow in large-scale systems (Keefer and Caudill, 2014; Neville et al., 2006). Such forms of pre-mating isolation reduce the risk of producing offspring with intermediate phenotypes that may be maladapted to both parental environments (Mendelson and Safran, 2021) and can be an important factor in maintaining local adaptation (Servedio and Boughman, 2017) and sometimes promoting incipient speciation (Shaw et al., 2024).

Salmonid homing behavior has long been recognized as a key driver of population structuring across river systems (Keefer and Caudill, 2014; Neville et al., 2006). This behavior is thought to arise primarily from olfactory imprinting during early development, allowing returning adults to recognize chemical cues from their home rivers (Dittman and Quinn, 1996; Hasler and Scholz, 2012; McCormick et al., 1998; Nordeng, 1977). Moreover, evidence from other anadromous fishes indicates additional reliance on visual landmarks and compass-based orientation (McCleave, 1967; Salmenkova, 2017; Stabell, 1984). While it is well known that this homing behavior can create population structuring among rivers (King & Stevens, 2021), it may also contribute to genetic differentiation among tributaries of the same river. In very large river systems (as defined by Munné & Prat, 2004), such as the Columbia and Fraser rivers (catchment area ∼ 670,000 km^2^ and 230,000 km^2^, respectively), extensive research using microsatellite and Single Nucleotide Polymorphism (SNP) markers has revealed pronounced genetic differentiation among salmonid populations, for example in the Pacific Chinook salmon (*Oncorhynchus tshawytscha*), inhabiting distinct tributaries (Christensen et al., 2024; Hess et al., 2016). These genetic breaks are often aligned with environmental gradients, suggesting that homing fidelity interacts with local adaptation to maintain population boundaries. For instance, populations spawning in cold, fast-flowing headwater streams tend to diverge from those spawning in slower, warmer, lowland tributaries, with differences linked to selection pressures such as thermal regimes, flow variability, and substrate composition (Fraser et al., 2011; Keefer and Caudill, 2014).

Studies of Atlantic salmon (*Salmo salar*) and brown trout (*Salmo trutta*) in the large river systems Varzuga (northwest Russia), Miramichi and St. John (eastern Canada), have demonstrated significant genetic divergence among populations inhabiting neighboring tributaries or even within single river networks (catchment area ∼ 9,000 and 54,000 km^2^) (Primmer et al., 2006; Verspoor et al., 2005). Similarly, sockeye salmon (*Oncorhynchus nerka*) in Lake Washington (United states) exhibit distinct genetic and phenotypic differences between stream and beach-spawning populations within the same watershed (Hendry et al., 2000), reflecting adaptation to contrasting spawning habitats. High-resolution genomic approaches, such as Restriction-site Associated DNA sequencing (RADseq), have revealed that such structuring can persist despite the absence of obvious physical barriers, suggesting a strong role for behavioral mechanisms like homing and assortative mating (Hecht et al., 2013). The interplay between behavioral isolation via homing and ecological selection across microhabitats likely acts synergistically to reinforce genetic structuring at multiple spatial scales. Such processes not only limit gene flow between conspecific populations but may also contribute to incipient reproductive isolation within species (Fraser et al., 2011; Quinn, 2018). These findings underscore the importance of considering both large-scale and fine-scale behavioral and ecological factors when assessing population structure in salmonids and other anadromous fishes.

The river Rönne å catchment in Scania, southern Sweden, sustains an Atlantic salmon population currently classified as Near Threatened on the IUCN Red List at the global scale due to significant population declines in recent years (IUCN, 2022). Salmon stocks have been identified as being in urgent need of targeted recovery actions to reach sustainable levels (Mannerla et al., 2011). By focusing on this smaller (catchment area 1,894 km^2^), human-impacted river system, and using high-resolution RADseq data from salmon fin clips collected across the tributaries of the Rönne å river, we aim to evaluate whether behavioral isolation, in the form of homing behavior, can lead to genetic differentiation in this small-scale river system. Understanding such patterns is crucial for informing conservation and management strategies in fragmented river systems, where preserving genetic diversity and local adaptation is key to supporting resilient salmon populations under ongoing anthropogenic pressures.

## Materials and methods

### Sample collection

A total of 90 anal fin clips from wild juvenile 0+ and 1+ of Atlantic salmon (7-168 mm) were collected across four tributaries within the Rönne å river system in Scania, southernmost Sweden: Klövabäcken (N=29), Rössjöholmsån (N=20), Bäljane å (N=21), and Pinnån (N=20) (Figure. 1, Supplementary table 1). The Rönne å river system extends approximately 85 km; however, Atlantic salmon are confined to the lower 38 km due to three mainstem hydropower dams that fragment the river and act as complete barriers to upstream movement. The total catchment area of the river covers about 1,894 km², of which the accessible sections downstream of the dams span 969 km². Field sampling was performed from August 2020 to May 2022 through electrofishing according to Bohlin et al. (1989), and individual tissue samples were collected by cutting a piece of the anal fin of caught individuals. The fin clips were conserved in 95% ethanol and stored in room temperature until DNA extraction.

**Figure 1.**
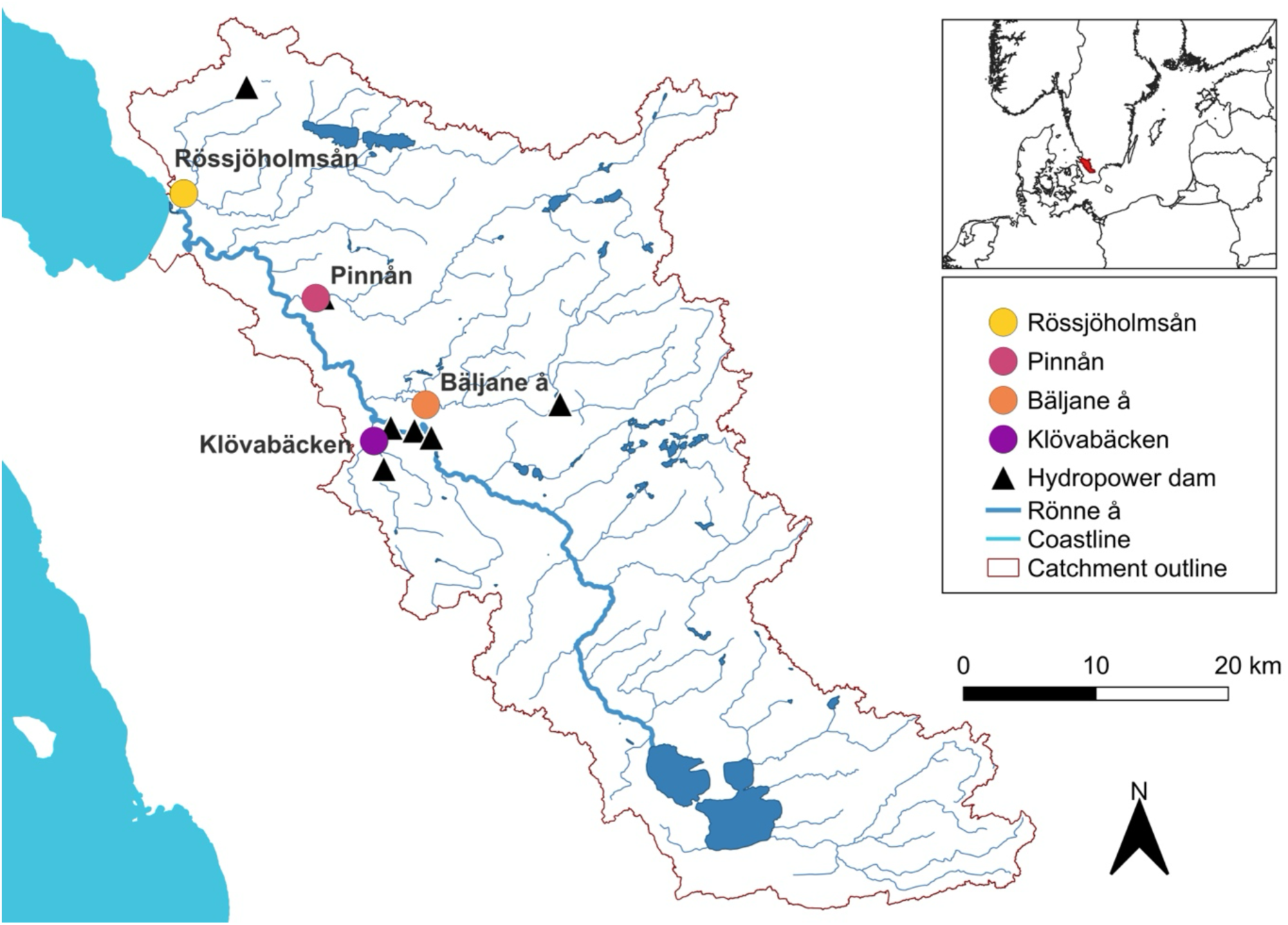
Map showing the sampling locations for Atlantic salmon (*Salmo salar*) in the river Rönne å catchment area. Dams that form complete migration barriers are represented by triangles. The orange line represents the river Rönne å catchment area.

### DNA extraction

Up to 25 mg of tissue was used to extract genomic DNA from each fin clip using Qiagen DNeasy Blood and Tissue Kit^®^ (Qiagen, Germany). To improve the extraction yield and obtain RNA-free DNA, 25 μl Qiagen Proteinase K and 4 μl Qiagen RNAse (100 mg/ml) were used, and the genomic DNA was eluted twice in 100 μl of pre-heated (60°C) Elution buffer (EB, Qiagen, Germany). Following the extraction, the concentration of DNA was measured in each sample with dsDNA BR Qubit Quant-iT assay (Invitrogen, USA). All samples below 20 ng/μl were four-fold concentrated, using the Genomic DNA Clean & Concentrator^TM^-10 Kit (Zymo Research, USA).

### RAD libraries preparation

The RAD library preparation protocol was modified and adapted from Amores et al., 2011 and Etter et al., 2011. Prior to processing, all DNA samples were diluted in EB to a concentration of 25 ng/μl, to minimize sequencing bias. The genomic DNA was then digested using the *SbfI-HF* restriction enzyme (New England Biolabs, USA) and bound to a P1 adapter using T4 DNA ligase (New England Biolabs, USA).

Subsequently, RAD libraries were constructed using pools of indexed individuals. Each pooled sample was sheared, using a focused ultrasonicator (M220 with XTU insert, Covaris), choosing an optimal length of 450 bp. All the fragments were cleaned and size-selected to a range of 300-600 bp, using Agencourt^®^ AMPure^®^ XP Beads (Beckman Coulter, USA). For left-side size selection, a ratio of beads to sample of 0.775 was used. Fragments longer than 600bp were removed using a ratio of 0.575 beads to sample.

After size-selection, the ends of the fragments were blunted using Blunt Enzyme Mix (New England Biolabs, USA) and bound to a 3’-dA overhang with Klenow fragment (3’è5’ exo-, New England Biolabs, USA). P2 adapters were added to each pool using T4 DNA ligase (New England Biolabs, USA). The PCR was then performed with 16 μl of genomic DNA, 4 μl of Solexa forward primer, 4 μl of Solexa reverse primer, 50 μl Phusion HF Master Mix (New England Biolabs, USA) and 38 μl PCR-grade water at 98°C initial denaturation for 30 s, followed by 18 cycles of 98°C for 10 s, 65°C for 30 s, and 72°C for 30 s, and a final extension at 72°C for 5 min. After purifying the product from residual primers using 1.0 beads to sample ratio, they were measured using dsDNA HS Qubit Quant-iT assay (Invitrogen, USA), to check that each pool had a concentration higher than 3 ng/μl. The pools were proportionally mixed and sent to the SNP&SEQ Technology Platform of the SciLifeLab facility in Uppsala, Sweden, where a paired-end sequencing with a read length of 150 bp was performed on an Illumina Nova Seq 6000 with 1.5 sequencing chemistry and 10% PhiX spike-in. The samples were sequenced in five libraries with included salmon samples from other projects not related to Rönne å.

### Data processing and analysis

All the sequences from salmon RAD libraries were processed using the software Stacks 2 v2.62 (Rochette et al., 2019). The Stacks function process_radtags was used for demultiplexing the samples based on inline barcodes (allowing for one mismatching base in the barcode), and filtering reads. All the reads identified as low-quality reads, with uncalled bases, no intact *SbfI-HF* restriction enzyme cutsite, and with adapter sequences (allowing for two mismatched bases) were removed. Additional quality checks were performed using FastQC v0.11.9 and MultiQC v1.11 (Ewels et al., 2016). The resulting loci were compared and aligned to a reference genome (Ssal_v3.1; GCF_000233375.1), using the function ref_map.pl. All the reads with a coverage below 2.874x (i.e., median coverage) were not included in further analysis. The populations function was subsequently run considering only the first SNP of each locus, to avoid reading multiple linked close SNPs. The setup for the populations run was p = 4 and r = 0.8, where p represents the number of populations and r is the minimum percentage of individuals in each population a locus must be present in. The output included a variant call format (VCF) file, which was used in subsequent analyses.

The calculation of the population pairwise F_ST_ values was performed in GenoDive v 3.06, with significance determined by 999 permutations (Osmond et al., 2023). Average nucleotide diversity within populations (π) and average absolute genetic divergence between populations (dxy) were estimated with Pixy v2.2.1 (Korunes and Samuk, 2021), using 100-kb non-overlapping genomic windows on the VCF file that had both variant and invariant sites. Discriminant Analysis of Principal Components (DAPC) of the individuals was realized using the R (v4.5.1) package adegenet, using the optim.a.score function to determine the optimal number of principal components to retain in the DAPC (Jombart, 2008).

Variant data were converted from VCF to BED, BIM, and FAM format using PLINK v1.9 (Purcell et al., 2007), and filtered to exclude SNPs with a minor allele frequency < 0.05 and missing genotype rate > 10%. Admixture analyses were then conducted in the Admixture v1.3.0 software (Alexander et al., 2009), with K values ranging from 1 to 4 using the cross-validation (-cv) procedure to determine the optimal number of clusters. The K value with the lowest cross-validation error was selected. Individual ancestry proportions were visualized as bar plots in R (v4.5.2) package tidyverse and ggplot2.

### Geographical data elaboration

QGIS v3.16 Hannover was used for the elaboration of all geographical data. All the coordinates were registered at the sampling site, and the maps were provided by HOT (Humanitarian OpenStreetMap Team, https://data.humdata.org/dataset/hotosm_swe_ waterways) and Orogènesis Soluciones Geograficas (http://tapiquen-sig.jimdo.com).

## Results

The total number of sequenced reads in the 5 salmon libraries (including the 90 samples included in this study but also samples from other projects) was 5,595,791,302, of which 4,921,974,195 (87.96%) were retained after demultiplexing and filtering. The mean number of sequenced reads per library was 1,119,158,260 (S.D. 201,117,754), of which an average of 984,394,839 (S.D. 171,079,970) was retained. A mean number of 79,356 loci was assembled in all samples. After filtering, 40 high-quality samples were retained, with an average coverage of 6.330x (ranging from 2.914x to 14.515x). A total of 709,708 loci were contained in all reads from the 40 samples that remained after the filtration and selection steps. Using the Stacks software, individuals were analyzed at the population level, resulting in a catalog of 71,312 shared loci, of which 21,301 were identified as variant loci.

### Population differentiation

The admixture analysis of the salmon SNP data supported a K=1 as the most likely number of clusters, suggesting that there are no deep ancestral clusters or deeply diverged clusters that split the population (Figure 2, Supplementary figure 1).

**Figure 2.**
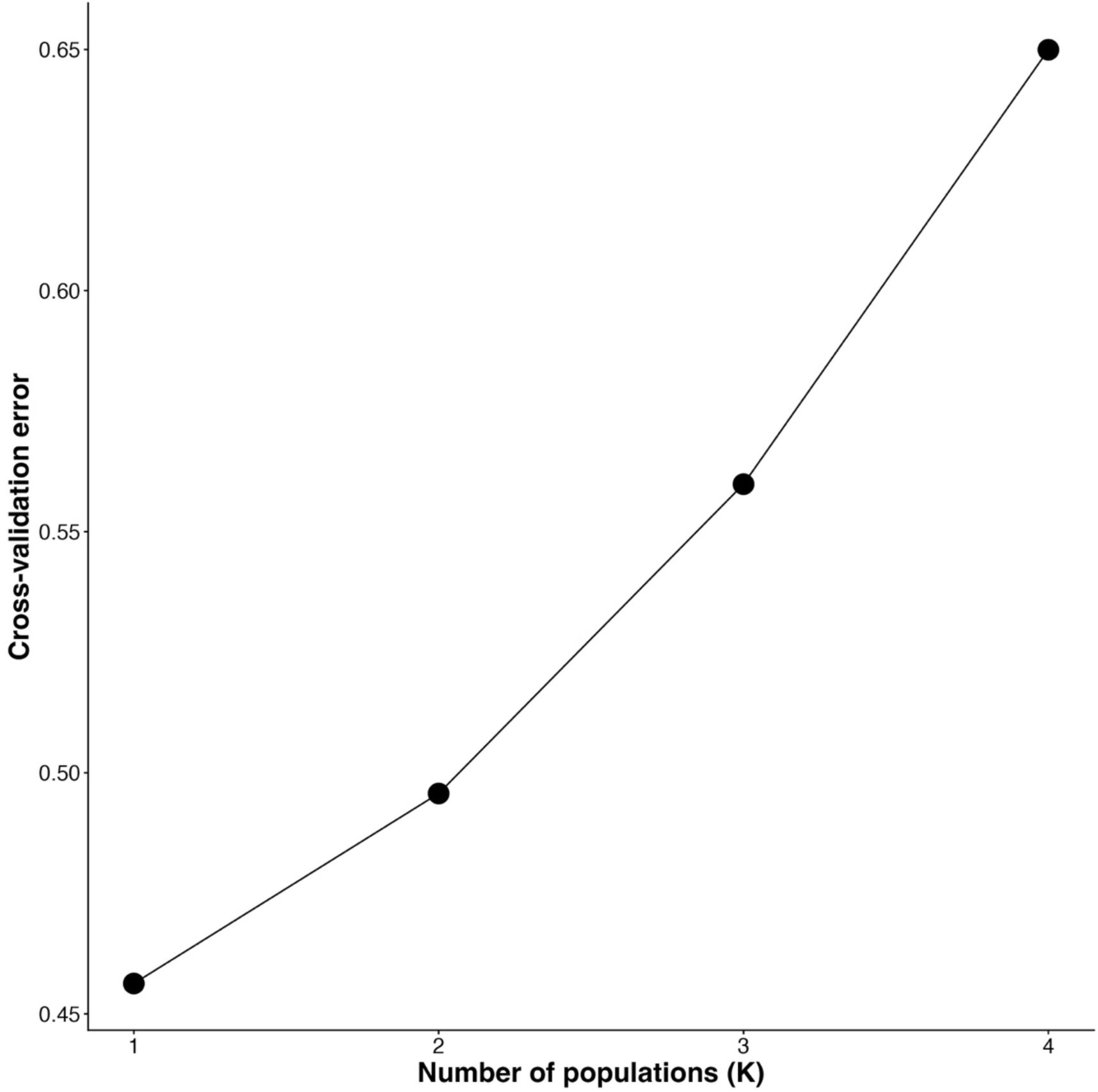
Cross-validation (CV) error values from ADMIXTURE analyses across K values (K = 1–4), showing the lowest CV error at K = 1. Structure plots for all K are available as supplementary figure 1.

The nucleotide diversity (π) was similar among the four populations. The highest nucleotide diversity was observed in Pinnån (π = 0.00186), closely followed by Klövabäcken (π = 0.00184), whereas Bäljane å (π = 0.00145) and Rössjöholmsån (π = 0.00150) exhibited slightly lower levels.

Pairwise absolute genetic divergence (Dxy) was consistently low, ranging from 0.00159 between Bäljane å and Rössjöholmsån to 0.00198 between Klövabäcken and Pinnån (Table 1). However, the pairwise F_ST_ values and DAPC revealed genetic differences among most sampling locations. Specifically, the calculated pairwise F_ST_ values were significant in all but three cases, ranging from 0.00 to 0.044 (Table 1, Supplementary Table 2). Non-significant F_ST_ values only appeared in comparisons where the individuals from Bäljane å were present. With these exceptions, all tributaries in the catchment area showed significant pairwise genetic differences, with the highest F_ST_ between Rössjöholmsån and Klövabäcken (0.044) (Table 1).

**Table 1.** Pairwise F_ST_ and dxy values for the analysis with four populations. Significant F_ST_ values (p<0.05) are represented by the * symbol. Lower triangle F_ST_. Upper triangle dxy.

| $F_{ST}$ values | Klövbäcken | Pinnån | Bäljane å | Rössjöholmsån |
| --- | --- | --- | --- | --- |
| Klövbäcken | - | 0.00198 | 0.00180 | 0.00178 |
| Pinnån | 0.016* | - | 0.00176 | 0.00176 |
| Bäljane å | 0.011 | 0.00 | - | 0.00159 |
| Rössjöholmsån | 0.044* | 0.037* | 0.037* | - |

Moreover, the DAPC of the four populations revealed differentiation of the population clusters from Klövabäcken and Rössjöholmsån. Whereas the Pinnån and Bäljane å individuals were closely clustered (Figure 3).

**Figure 3.**
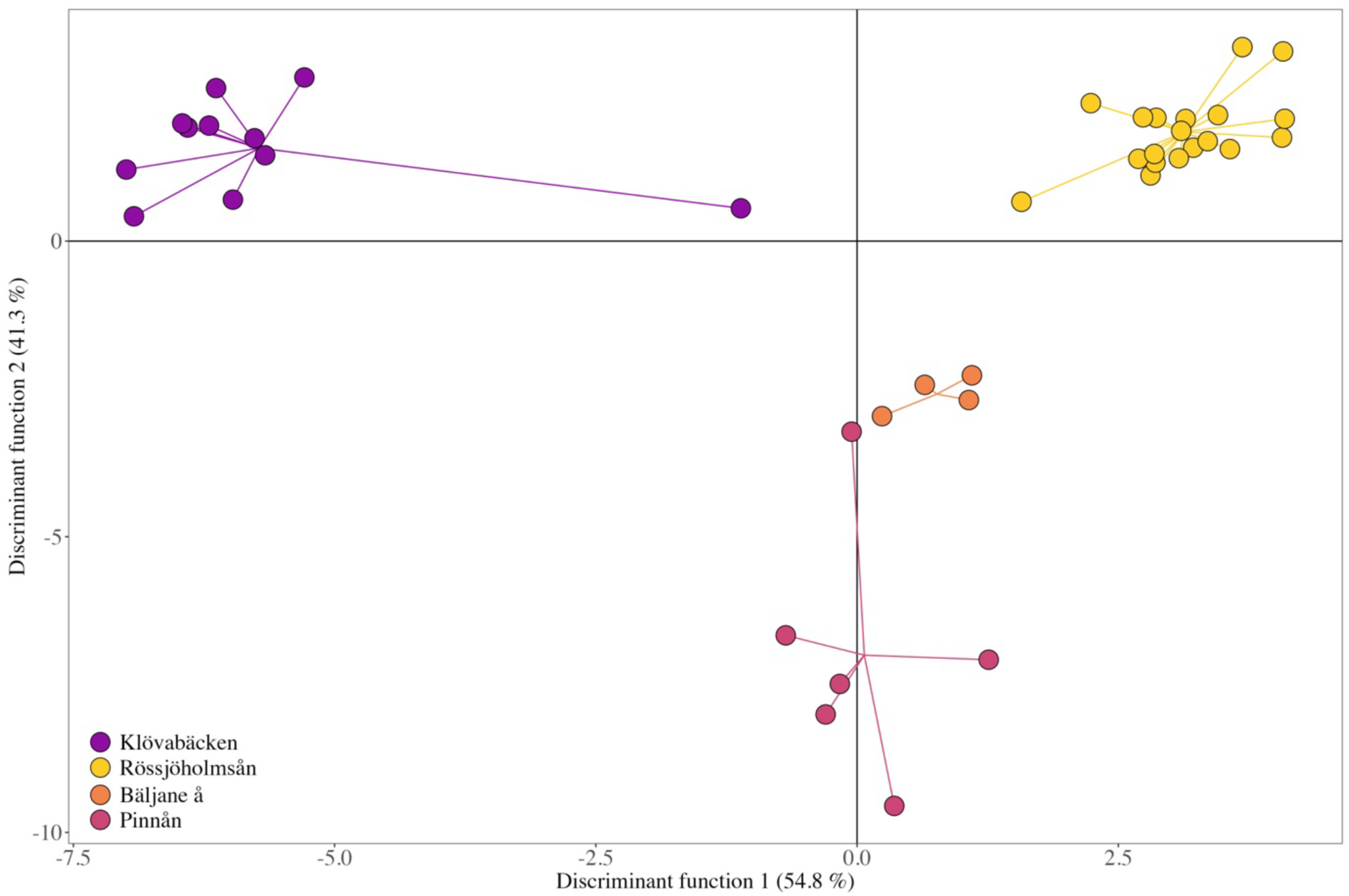
DAPC analysis of the four different salmon populations. Discriminant function 1 is represented by the x-axis and discriminant function 2 is represented by the y-axis. Klövabäcken n=11, Rössjöholmsån n= 19, Bäljane å n= 4, Pinnån n=6.

## Discussion

Understanding fine-scale genetic population structure within wild Atlantic salmon watersheds is critical for effective conservation and management, particularly given the increasing pressures from habitat fragmentation, climate change, and anthropogenic impacts. While it is well known that salmon from separate rivers form distinct populations (King et al., 2001), evidence for population structure within large and medium-sized rivers has also been found in salmonids (Christensen et al., 2024; Hess et al., 2016; Primmer et al., 2006; Verspoor et al., 2005), highlighting the possibility that a single large river system may have to be subdivided into separate management units. However, it is not known if smaller river systems can also harbor genetically distinct salmon populations. This study provides novel insights into the fine-scale genetic differentiation among salmon populations in a small river system, emphasizing the role of natal homing behavior and landscape features in shaping population boundaries within this small river.

Despite the absence of clearly defined genetic clusters inferred by the Admixture analysis for Rönne å catchment (Figure 2, Supplementary Figure 1), the presence of significant pairwise F_ST_ values and DAPC demonstrates that population differentiation is emerging within the river system (Figure 3, Table 1). This interpretation is further supported by the similar and low nucleotide diversity (π) among the populations and the low Dxy values across populations, suggesting limited absolute sequence divergence and consistent recent or incipient divergence. Such scenarios are expected in systems where some gene flow is still ongoing, and the different populations share the same historical ancestors, preventing strong ancestral structure despite localized reproductive structuring (Jones and Wang, 2012; Nutt, 2008). However, continued sampling in the area is essential to determine whether these patterns remain consistent with an increase in the number of individuals analyzed.

The Admixture supported K = 1 as the most likely number of ancestral clusters; however, exploratory analyses at higher K values (K = 2–4) revealed a tendency for individuals from Klövabäcken and Rössjöholmsån tributaries to show partial differentiation from the remaining sampling locations. This pattern is consistent with the significant pairwise F_ST_ values and the clustering observed in DAPC analyses. In larger river systems, different genetic clusters have been identified using Admixture/Structure, e.g. in the Fraser River for Pacific salmon (Christensen et al., 2024), as well as the Teno River for Atlantic salmon (Vähä et al., 2007), likely reflecting stronger genetic differences between subpopulations. For example, in the Teno River, pairwise F_ST_ ranged between 0.015 to 0.201 for the inferred populations, with an average of 0.092 (Vähä et al., 2007), while we estimated the maximum pairwise F_ST_ in the Rönne å to be substantially smaller at 0.045. Nevertheless, studies of selection in natural populations generally find strong divergent selection between populations even at F_ST_ < 0.05 (Leinonen et al., 2013), and although we did not measure selection on traits and cannot assess whether the population differentiation is caused by selection or drift, local adaptation is often detected within our range of population differentiation (Leinonen et al., 2013, 2008).

Considering the pairwise F_ST_ values and the clustering observed in DAPC analyses, Klövabäcken and Rössjöholmsån tributaries formed distinct populations (Figure 3, Table 1), suggesting the emergence of genetic differentiation within the Rönne å system. Klövabäcken and Rössjöholmsån exhibit distinct topographical and hydrological features compared to the other affluents, and also are the two furthest tributaries from each other (approximately 30 km apart). This geographical distance may have contributed to reduced gene flow between the two populations. Rössjöholmsån is the first tributary encountered by upstream-migrating salmon reentering the watershed, which could enhance its recognition as a return site and explain the marked genetic differentiation between the Rössjöholmsån population and those from other locations (Figure 3). Conversely, Klövabäcken is the only accessible tributary draining from the south, and together with its small catchment area and lower discharge (Supplementary table 3) could likely increase the influence of genetic drift.

In addition, pairwise F_ST_ values and DAPC analyses revealed that the population in Pinnån clustered distinctly from all other locations, with the exception of Bäljane å (Figure 3, Table 1). The closer clustering of Pinnån and Bäljane å individuals may be attributed to their geographic proximity and similar environmental characteristics, including similar catchment sizes, discharge ranges, mean water temperatures, as well as forest cover, which may contribute to analogous ecological conditions (Supplementary table 3). Both tributaries enter the river system from the northern side of the catchment and lie only a few kilometers apart, potentially increasing opportunities for straying between them (Figure 1). Such spatial closeness can facilitate gene flow if some individuals home to the “wrong” tributary, a phenomenon documented in other salmonid populations (Keefer and Caudill, 2014; Quinn, 1993). However, this interpretation should be treated with caution, as the observed clustering could also be influenced by the smaller sample size, potentially reducing the power to detect differentiation. Nevertheless, their position in the DAPC suggests that these two groups may form a single genetic population that is distinct from all other sampled populations in the watershed (Figure 3).

Overall, the findings of this study demonstrate evidence of fine-scale genetic structuring within the Rönne å watershed, influenced by environmental, geographic, and behavioral factors, consistent with previous research showing that local adaptation, homing fidelity, and habitat diversity promote genetic differentiation even at small spatial scales (Kitanishi et al., 2009; Osmond et al., 2023). However, it is important to acknowledge that low sample sizes and the presence of closely related individuals can potentially inflate estimates of population structure (Manunza et al., 2025), and thus further investigation is required to fully elucidate the genetic structuring within the Rönne å catchment and the precise mechanisms governing within-river differentiations. The presence of multiple genetically distinct populations underscores the importance of conserving this diversity, especially considering historical anthropogenic impacts such as hydropower development that have contributed to genetic homogenization over the past century (Östergren et al., 2021).

## Conclusions

Our results reveal emergent structuration among Atlantic salmon populations within the Rönne å river system, with signals of subtle differentiation between populations in different tributaries. This pattern suggests that, despite longitudinal connectivity, gene flow is not uniform and is likely shaped by a combination of variations in dispersal, spawning homing behavior, and site characteristics. From a conservation and fisheries management perspective, these findings highlight the importance of recognizing the particularities of the independent subpopulation units rather than considering them as a single, well-mixed population. Management strategies should therefore aim to preserve connectivity while also protecting local habitats and spawning areas, particularly in smaller tributaries that may harbor unique genetic variation. Maintaining habitat heterogeneity is essential to sustain both local adaptation and population resilience in the face of environmental change.

## Acknowledgements

We thank Karin Rengefors for supporting the development of the RAD sequencing protocol, and for providing helpful comments in the application for this research, laboratory tips, and improvement of the manuscript. The DNA laboratory in Lund, especially for Marie Svensson, for providing infrastructure and support for molecular analyses. SciLifeLab and the National Genomic Infrastructure for assistance with massive parallel sequencing.

## Data availability

The datasets generated during and/or analysed during the current study are available from the corresponding author upon request.

## Author contributions

Conceptualization: JS, AP, OC, PAN

Data curation: FDG, COC, JS, ML, RG, AP, PAN

Formal analysis: FDG, COC, JS, ML, RG

Funding acquisition: JS, ML, AP, OC, PAN

Investigation: FDG, COC, JS, AP, OC, SS, PAN

Methodology: FDG, COC, JS, ML, RG, AP, OC, SS, PAN

Resources: JS, AP, OC, SS, PAN

Supervision: JS, RG, PAN

Visualization: FDG, COC, ML

Original draft: FDG, COC, JS, ML, AP, PAN

Review and editing: FDG, COC, JS, ML, RG, AP, OC, SS, PAN

## SUPPLEMENTARY MATERIAL

**Supplementary Figure 1.**
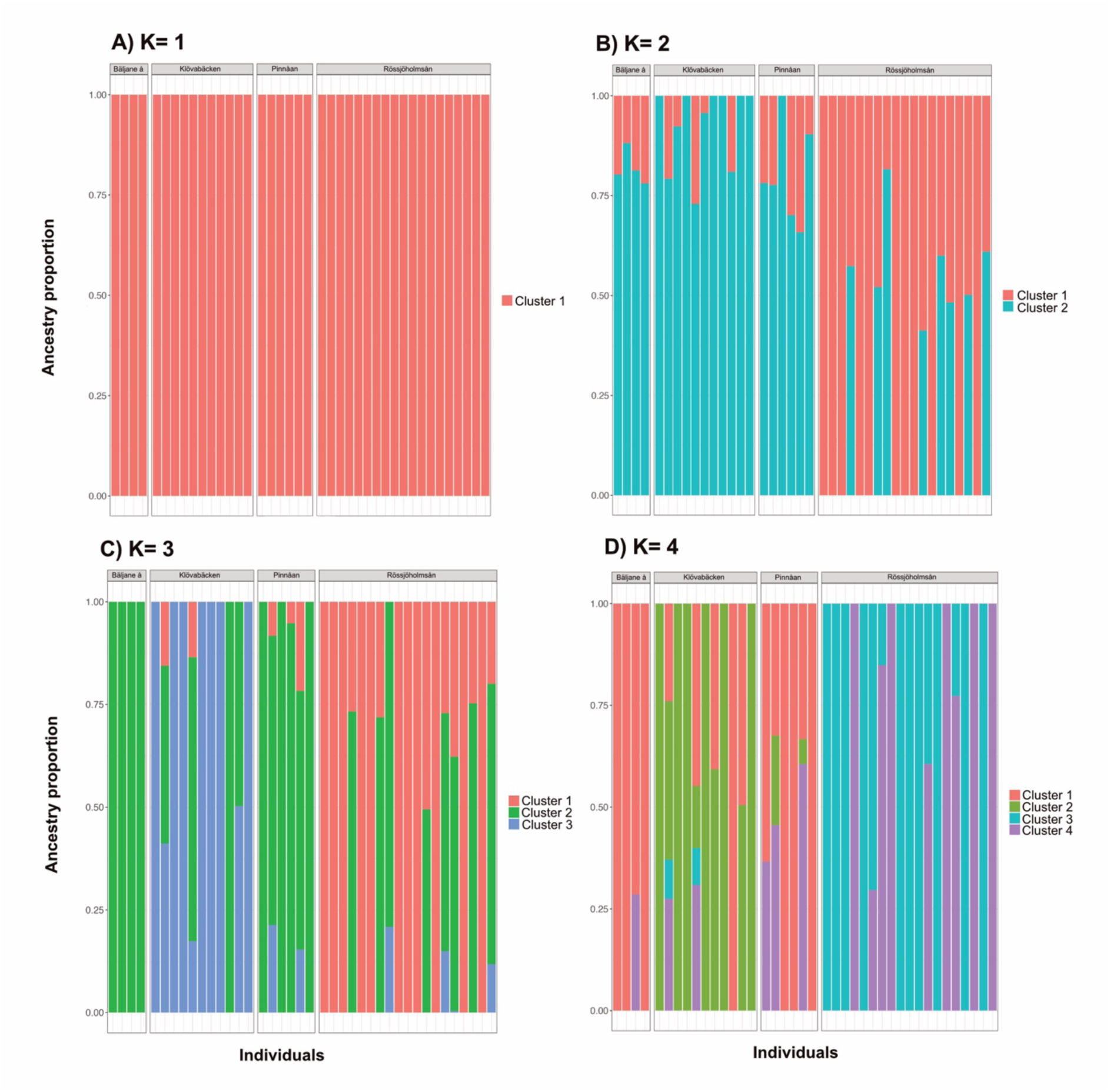
**ADMIXTURE clustering results for K 1–4 based on the assumption of four different population datasets for Ronne å.**

**Supplementary Table 1.** Overview of sampling location, GPS coordinates, number of fin clips collected, and number of retained individuals.

| Species | Area | Total fin clips samples | Retained individuals | Coordinate X (east) | Coordinate Y (north) |
| --- | --- | --- | --- | --- | --- |
| *S. salar* | Pinnån | 20 | 6 | 13.019881 | 56.209620 |
|  | Klövbäcken | 29 | 11 | 13.095639 | 56.113864 |
|  | Rössjöholmsån | 20 | 19 | 12.855185 | 56.277816 |
|  | Bäljane å | 21 | 4 | 13.157246 | 56.139370 |

**Supplementary Table 2.**
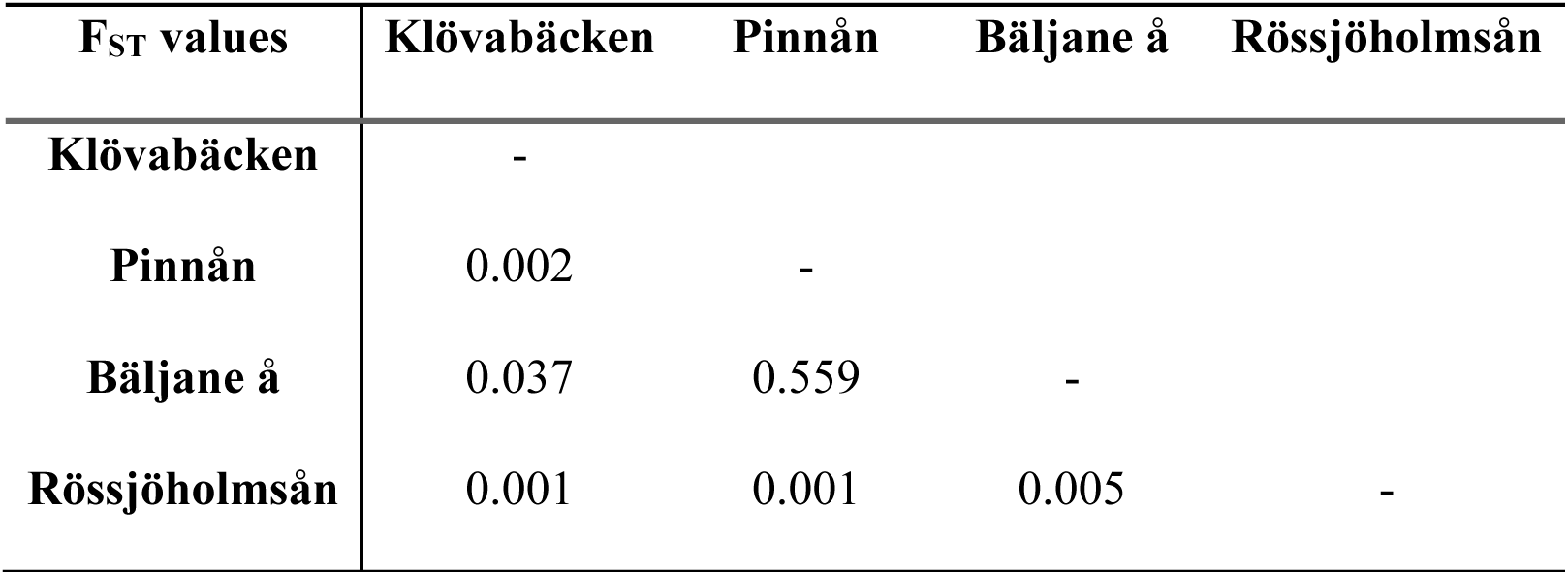
P-values of the Pairwise F_ST_ analysis for the populations studied.

**Supplementary Table 3.** River and catchment characteristics of Rönne å and its tributaries. Discharge (MQ) and water temperature (T) are annual means and coefficients of variation (C.V.) of daily estimates 2020-2022 (SMHI, 2026).

| **River Unit** | **Catchment** | **MQ (C.V.)** | **T (C.V.)** | **Forest** | **Agriculture** | **Lakes** |
| --- | --- | --- | --- | --- | --- | --- |
|  | **km<sup>2</sup>** | **m<sup>3</sup>s<sup>-1</sup>(%)** | **°C (%)** | **%** | **%** | **%** |
| **Rönne å** | 1302 | 12.6 (104) | 9.7 (64) | 41 | 30 | 4.6 |
| **Rössjöholmsån** | 263 | 3.6 (107) | 9.7 (64) | 39 | 31 | 4.4 |
| **Pinnån** | 211 | 2.1 (105) | 9.4 (67) | 52 | 47 | 1.5 |
| **Bäljane å** | 241 | 3.2 (123) | 9.6 (66) | 50 | 17 | 1.0 |
| **Klövbäcken** | 47 | 0.48 (133) | 9.2 (65) | 49 | 32 | 1.2 |

Reference: https://vattenwebb.smhi.se/modelarea/. Uploaded 202601

